# Loss of a conserved MAPK causes catastrophic failure in assembly of a specialized cilium-like structure in *Toxoplasma gondii*

**DOI:** 10.1101/2020.02.02.931022

**Authors:** William J. O’Shaughnessy, Xiaoyu Hu, Tsebaot Beraki, Matthew McDougal, Michael L. Reese

## Abstract

Primary cilia are important organizing centers that control diverse cellular processes. Apicomplexan parasites like *Toxoplasma gondii* have a specialized cilium-like structure called the conoid that organizes the secretory and invasion machinery critical for the parasites’ lifestyle. The proteins that initiate the biogenesis of this structure are largely unknown. We identified the *Toxoplasma* ortholog of the conserved kinase ERK7 as essential to conoid assembly. Parasites in which ERK7 has been depleted lose their conoids late during maturation and are immotile and thus unable to invade new host cells. This is the most severe phenotype to conoid biogenesis yet reported, and is made more striking by the fact that ERK7 is not a conoid protein, as it localizes just basal to the structure. ERK7 has been recently implicated in ciliogenesis in metazoan cells, and our data suggest that this kinase has an ancient and central role in regulating ciliogenesis throughout Eukaryota.

## Introduction

Cilia are specialized organelles that mediate cell motility and signal transduction. In apicomplexan parasites, the cilium was likely adapted to form the organizing core of the “apical complex” of cytoskeletal structures and secretory organelles for which the phylum is named (Figure 1; de Leon et al., 2013). The protein components of the apical complex include orthologs of cilium-associated proteins including centrins (Hu *et al*., 2006; Lentini *et al*., 2019), a SAS6 cartwheel protein (de Leon *et al*., 2013; Lévêque *et al*., 2016), an associated rootlet fiber (Francia *et al*., 2012), and a distributed microtubule-organizing center called the apical polar ring (Leung *et al*., 2017). During the asexual stage, Apicomplexa such as *Toxoplasma* have a specialized cilium-like structure called the conoid, and typical motile flagella grow at this site during the sexual stage (Francia *et al*., 2015). As in ciliogenesis in model organisms, the *Toxoplasma* apical complex splits from the centrosome early during daughter cell budding (Anderson-White *et al*., 2012), but the molecular mechanisms that regulate this biogenesis are unknown.

**Figure 1:**
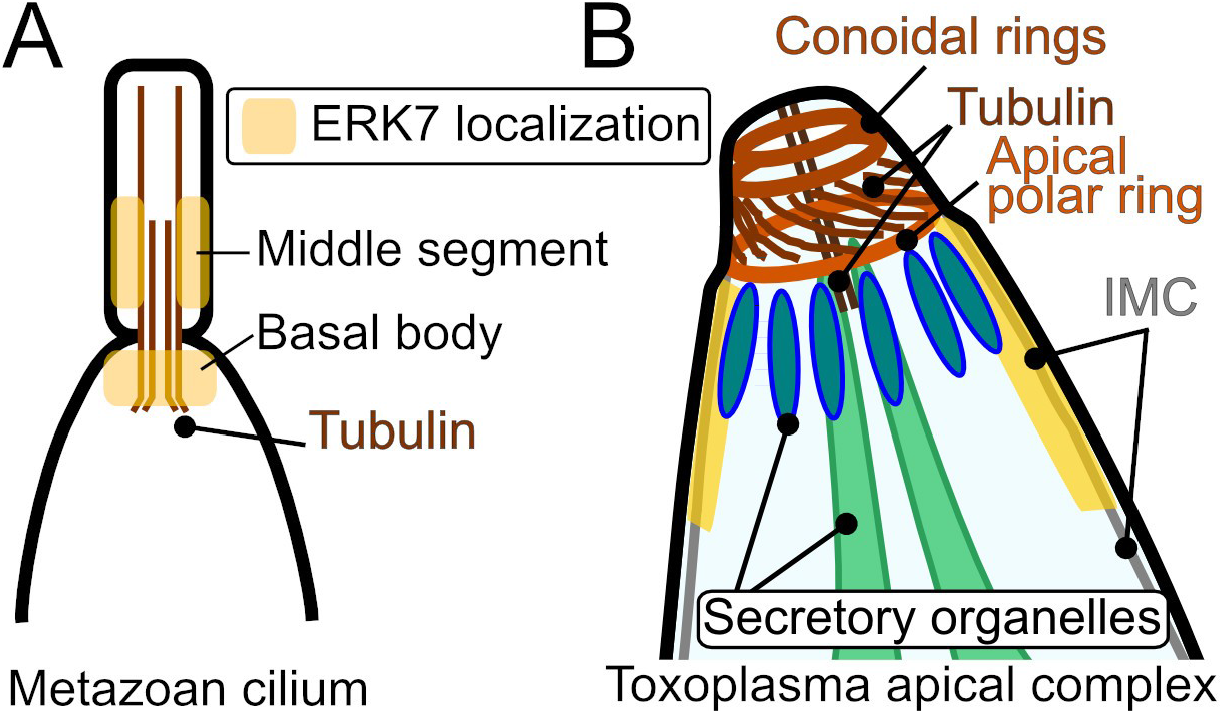
A cartoon comparing the (A) metazoa cilium and (B) *Toxoplasma* apical complex.

Here, we describe the essential role of the *Toxoplasma* ortholog of ERK7 in apical complex biogenesis. We found that ERK7 localizes to the apical end of the parasite throughout the cell cycle, suggesting a role in the parasite invasion machinery. Consistent with this hypothesis, we found that parasites lacking ERK7 protein have a complete block in egress and invasion. Without ERK7, parasites fail to develop a conoid, suggesting that ERK7 is required for its biogenesis. Even though ERK7 does not appear to be conoid-localized, its depletion causes complete loss of the conoid in mature parasites. Our findings are consistent with recent reports implicating metazoan ERK7 in ciliogenesis in invertebrate and vertebrate models (Miyatake *et al*., 2015; Kazatskaya *et al*., 2017), and suggest that the MAPK ERK7 is an essential component of the core machinery that regulates ciliogenesis throughout Eukaryota.

## Results and Discussion

### ERK7 is a conserved MAPK localized at the apical end of the parasite

The *Toxoplasma* genome encodes 3 predicted MAPKs. To assess whether the *Toxoplasma* gene TGME49_233010 is, indeed, a member of the ERK7 family, we estimated the phylogenetic tree from an alignment of ERK7 sequences from diverse organisms (Supplemental Figure S1), including other members of the CMGC kinase family as outgroups. The phylogenetic tree shows high bootstrap support for the TGME49_233010 as a member of this family.

To determine ERK7 localization within the parasite, we used CRISPR-mediated homologous repair to tag the endogenous locus with a C-terminal 3xHA epitope tag. Immunofluorescence analysis shows ERK7 signal concentrates at the apical end of the parasite. Specifically, ERK7 co-localizes with the apical cap protein ISP1, just basal to the tubulin-rich parasite conoid (Figure 2A). Furthermore, ERK7 appears to be recruited to this structure early in its biogenesis, as clear foci are evident in both mature parasites and in early daughter buds (Figure 2A). Structured illumination microscopy revealed punctate ERK7 staining throughout the parasites, and confirmed that ERK7 concentrates just basal to the apical complex ring in both mature and daughter parasites. Furthermore, at high resolution, ERK7 does not colocalize with either the parasite cortical microtubules (Figure 2B) or the apical cap cytoskeletal protein ISP1 (Figure 2C).

**Figure 2:**
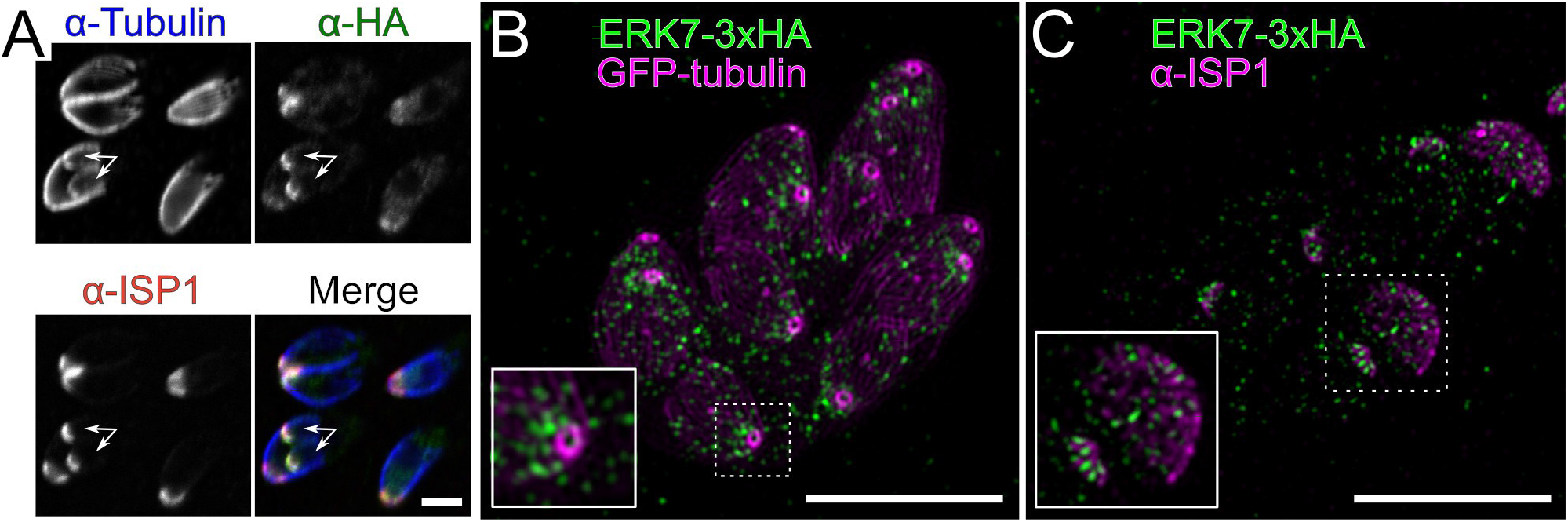
ERK7 is apically localized. (A) 0.5 μm confocal slice of immunofluorescence using antibodies for 3xHA-tagged ERK7 (green), β-tubulin (blue), and the apical cap marker ISP1 (red). Note that anti-tubulin does not stain the conoid, likely due to antigen accessibility. Arrows indicate daughter buds. Scale bar: 3 μm. Maximum intensity projection of SIM stacks of (B) ERK7-3xHA (green) and GFP-α-tubulin (magenta) and (C) ERK7-3xHA (green) and anti-ISP1. Scale bars: 5 μm.

### ERK7 is essential for parasite invasion and egress

We were unable to obtain ERK7 knockouts using either homologous recombination or CRISPR-mediated strategies. We therefore turned to the auxin-inducible degron (AID) system (Nishimura *et al*., 2009; Long *et al*., 2017b), which enables rapid, inducible degradation of a protein. We made a parasite strain in which the ERK7 protein was expressed in frame with an AID and 3xFLAG tag in the background of RH*Δku80* expressing the rice TIR1 auxin response protein (ERK7^AID^). ERK7 localization was unaffected by the AID tag (Supplemental Figure S2A). We found that ERK7^AID^ protein was rapidly degraded upon addition of the auxin indole-3-acetic acid (IAA), as it was undetectable by western blot 15 min after IAA treatment (Figure 3A). We will refer to parasites in which ERK7 has been inducibly degraded as ERK7^AID/IAA^. ERK7 was essential for the lytic cycle, as ERK7^AID/IAA^ produced no plaques (Figure 3B). To verify that this phenotype was specific to ERK7 depletion, we made parasites expressing a non-degradable copy of wild-type ERK7-3xHA in the background of the ERK7^AID^ parasites (Supplemental Figure S2B,C). As expected, wild-type ERK7 rescued the ability of the ERK7^AID/IAA^ parasites to form plaques (Figure 3B). However, we were unable to obtain stable parasites expressing kinase-dead (D136A) ERK7, suggesting expression of kinase-dead protein has a strong fitness cost.

**Figure 3:**
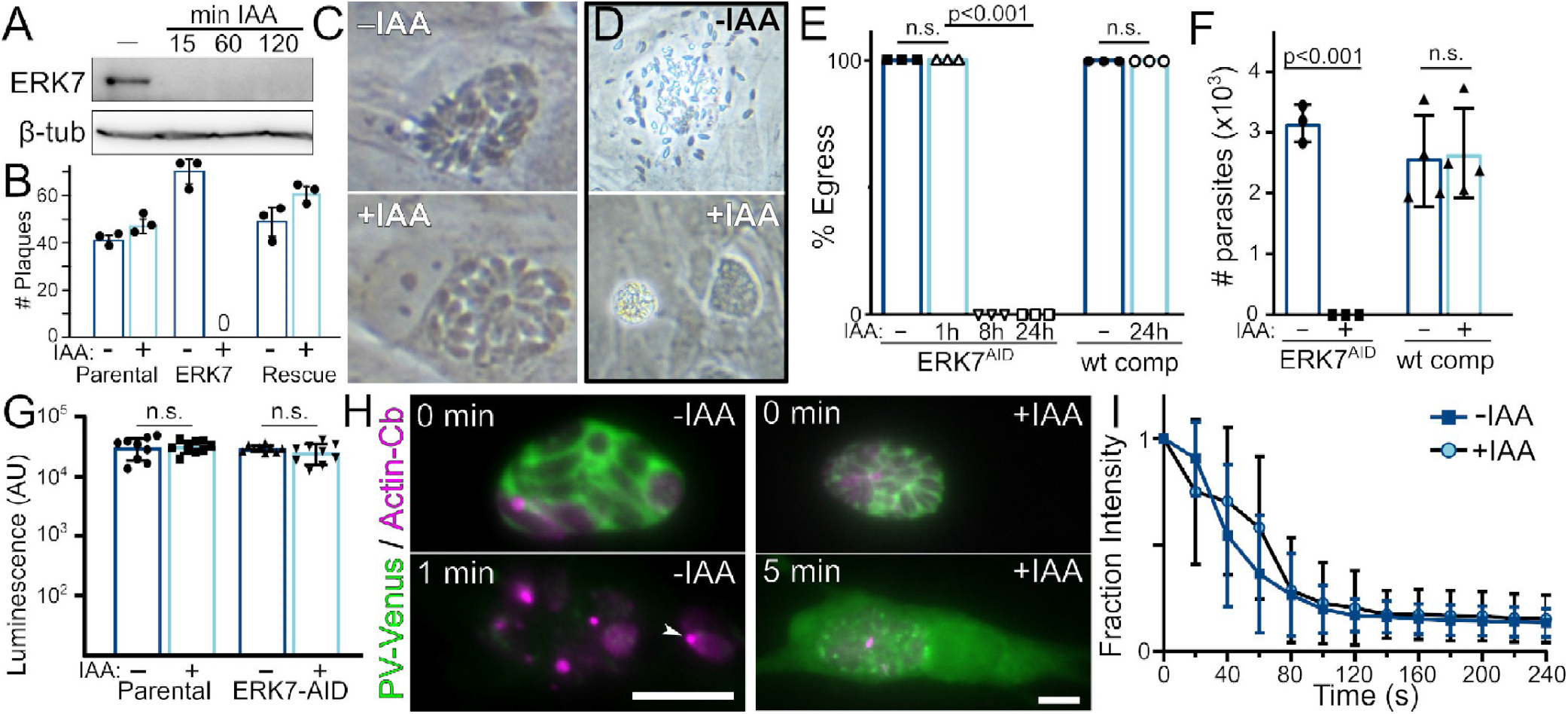
Loss of ERK7 blocks parasite invasion and egress. (A) ERK7^AID^ parasites were incubated as indicated with IAA and lysates were probed with anti-FLAG and anti β-tubulin as a loading control. (B) Quantification of 3 replicate plaque assays with parental, ERK7^AID^, and ERK7^AID^(wt-comp) strains +/- IAA. (C) Phase image of ERK7^AID^ parasites grown 24 h +/- IAA. (D) Parasites as in (C) were grown 18 h more until vacuoles began to rupture. (E) Quantification of egress from n=3 replicates grown as indicated in IAA. (F) Quantification of attached and invaded parasites from ERK7^AID^ and ERK7^AID^(wt-comp) strains grown +/- IAA. (G) Quantification of *Gaussia* luciferase activity secreted by the indicated strains grown +/- IAA. (H) ERK7^AID^ and ERK7^AID/IAA^ expressing an actin chromobody (magenta) and secreting Venus into the PV (green) were imaged at the indicated times after ionophore treatment. Scale bars: 10 μm. (I) Quantification of PV Venus signal of parasites as in (H) relative to first frame (n=93 -IAA; 81 +IAA). Error bars are s.d.

Failure to form plaques can be due to impairments in replication rate, invasion, or egress efficiency. While ERK7 has been suggested to be important for efficient replication (Li *et al*., 2016), we observed only a modest phenotype in ERK7^AID/IAA^ parasite replication (Figure 3C). This could not explain the lack of plaque formation. Instead, we noted that ERK7^AID/IAA^ parasites replicated until they mechanically disrupted the host cell. After host cell rupture, parasites appeared trapped within their vacuoles, which we found floating in the media (Figure 3D). Such trapped parasites suggest a block in natural egress reminiscent the deletion of parasite perforin-like protein 1 (PLP1) (Kafsack *et al*., 2009). Parasite egress from host cells can be induced by treatment with the calcium ionophore A23187 (Carruthers and Sibley, 1999). We therefore tested if ERK7 was essential for induced egress. Parasites were allowed to invade host fibroblasts for 4 h and then grown with or without IAA for 24 h. Egress was induced with ionophore. ERK7^AID/IAA^ parasites showed a complete block in egress (Figures 3E). Notably, when parasites were treated with a brief 1 h exposure to IAA, we observed no phenotype (Figure 3E), indicating that the effects we see require completion of a parasite cell cycle (which takes 6-8 h; Nishi et al., 2008). Consistent with this idea, parasites incubated for 8 h with IAA showed a similar block in egress to those treated for 24 h (Figure 3E).

We next tested whether parasites lacking ERK7 were able to invade host cells. ERK7^AID^ and rescued parasites were grown with or without IAA for 24 h. The parasites were mechanically released from the host cells and an equal number of each strain incubated with fresh host monolayers for 2 h at 37°C. Cells were then fixed and imaged. Strikingly, while ERK7^AID^ parasites faithfully attached and invaded the cells, we observed a complete block in both attachment and invasion in ERK7^AID/IAA^ parasites, which was rescued when complemented with a wild-type copy (Figure 3F). These severe egress and invasion phenotypes fully explain the null plaque count we observed (Figure 3B).

### ERK7 is required for parasite motility but not microneme secretion

Both egress and invasion require secretion of specialized organelles called the micronemes (Lourido *et al*., 2012). ERK7 localizes to the apical end of the parasite (Figure 2), near the site from which the micronemes are thought to be secreted. Thus, ERK7 could act as a regulator of secretion. To test this hypothesis, we engineered the parental and ERK7^AID^ strains to express MIC2 fused to *Gaussia* luciferase (Brown *et al*., 2016). Surprisingly, ERK7 depletion did not inhibit microneme secretion of this luciferase reporter (Figure 3G). The microneme protein PLP1 permeabilizes the parasitophorous vacuole (PV) to large macromolecules (Kafsack *et al*., 2009). To confirm that ERK7 depletion did not block secretion of native microneme proteins, we used strains that secreted mVenus into the PV and measured mVenus diffusion from the PV after ionophore treatment to confirm PLP1 secretion (Figure 3H, Supplemental Movie 1). For the majority of vacuoles imaged (>90%), we observed permeabilization of the PV membrane within the 5 min of imaging after ionophore treatment, independent of growth in IAA and with similar dynamics (Figure 3I). However, while ERK7^AID^ parasites lysed out of the host cell shortly after permeabilization, ERK7^AID/IAA^ parasites were immobile and trapped within their PV. For these ERK7^AID/IAA^ vacuoles, mVenus signal remained within the host cell, which was not itself permeabilized. Taken together, these data indicate that functional microneme secretion occurs in the ERK7^AID/IAA^ parasites, though they do not egress.

*Toxoplasma* parasites move using an unusual actin-based mechanism termed “gliding motility.” Actin nucleation at the parasite’s apical tip is essential to initiating this process (Tosetti *et al*., 2019). We used an established actin “chromobody” (Periz et al., 2017) to visualize F-actin dynamics during ionophore treatment. As expected, F-actin puncta quickly formed at the apical tips of the ERK7^AID^ parasites, preceding motility and the rapid egress described above (Figure 3H, Supplemental Movie 1). ERK7^AID/IAA^ parasites, however, showed no change in their actin structures over 5 min of imaging (Figure 3H, Supplemental Movie 1). Thus, parasites that have developed without ERK7 cannot regulate their actin structures to initiate movement.

### The parasite apical complex formation requires functional ERK7

The combination of phenotypes above and the requirement for a full cell cycle suggested that loss of ERK7 blocks development of structures critical to parasite motility, invasion, and egress. Recently, the *Toxoplasma* conoid has been implicated in mediating these processes independently of any role in microneme secretion (Long *et al*., 2017b, 2017a; Nagayasu *et al*., 2017; Tosetti *et al*., 2019). We therefore examined transmission electron micrographs (TEM) of intracellular ERK7^AID^ parasites that had been grown with or without IAA for 18 h. While we could easily identify the apical complex conoid with a “crown” of micronemes in ERK7^AID^ parasites (Figure 4A), we could find no evidence of a conoid in any ERK7^AID/IAA^ parasites (Figure 4B).

**Figure 4:**
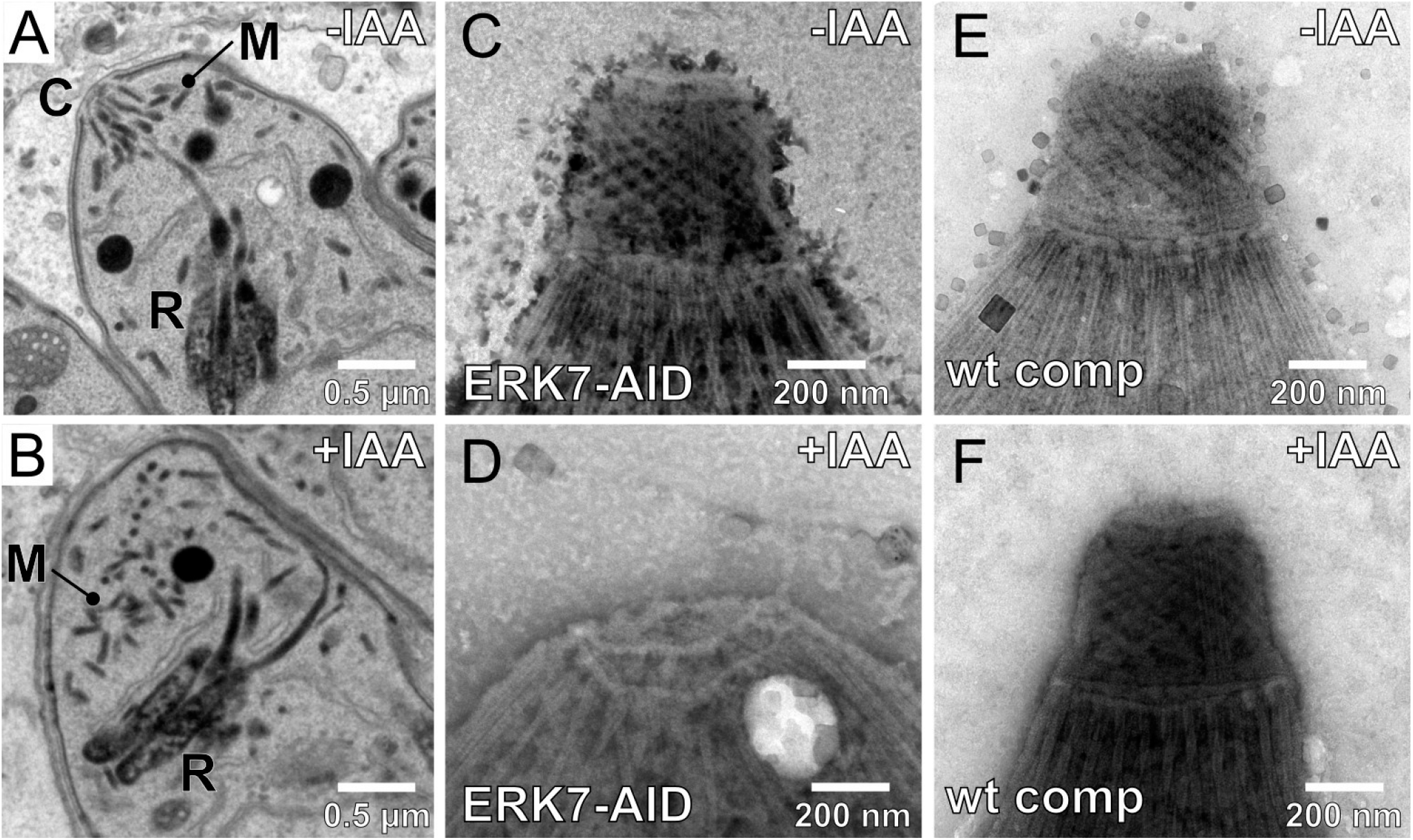
The apical complex is disrupted in parasites lacking ERK7. TEM of (A) ERK7^AID^ parasites develop normal conoids [C] and well organized secretory organelles (micronemes [M] and rhoptries [R]), while those grown (B) for 18 h in IAA have no identifiable conoid and disorganized secretory organelles. TEM of negatively stained detergent-extracted parasites reveal normal microtubule structures in ERK7^AID^ parasites (n=20) (C) including the distinctive rings and conoid spiral of the apical complex, which are lost when parasites are grown for 18 h in IAA (n=60) (D). Complementation of ERK7^AID^ with wild-type ERK7 rescues all phenotypes (n=20) (E,F).

The microtubule structures of the apical complex are preserved in detergent-extracted parasites. When we examined negatively stained detergent-extracted parasite “ghosts” by TEM (Figures 4C-F), we found that ERK7 depletion results in a complete loss of the conoid structure, which is rescued by complementation with wild-type ERK7. This is striking for two reasons. First, the complete loss of the conoid is much more severe than previously reported phenotypes that caused only its deformation (Long *et al*., 2017a; Nagayasu *et al*., 2017). Second, previously reported phenotypes in the apical complex have been caused by disruption of proteins that localize to the complex itself, and therefore likely perform a structural role. ERK7, however, is not an apical complex protein, as it localizes just basal to the structure (Figure 2). These observations suggest ERK7 performs a regulatory role in conoid biogenesis. Notably, this profound ultrastructural defect appears to be specific to the conoid, as the 22 cortical microtubules appear unaffected by ERK7 degradation (Figure 4D). Consistent with this, the apical polar ring that is thought to form the microtubule-organizing center for the cortical microtubules (Leung *et al*., 2017) appears to be preserved in parasites lacking ERK7.

In addition to the lack of conoids, we noted a striking disorganization of the micronemes and rhoptries in the TEMs of ERK7^AID/IAA^ parasites (Figure 4B). To confirm these observations, we imaged parasites either expressing fluorescent-protein fusions or stained with available antibodies to a variety of markers (Figure 5). Consistent with our TEM data, the apical secretory organelles appeared disorganized when visualized by fluorescence microscopy. Staining with RON11 (which marks the apical rhoptry neck) and ROP2 (which marks the basal rhoptry bulb), we found that rhoptries do not bundle together upon ERK7 degradation and are not consistently oriented with their necks towards the parasite apical end (Figure 5A). These data suggest that loss of the conoid causes a failure in rhoptry tethering.

Fluorescence microscopy also showed that the microneme marker PLP1 was not aligned at the parasite plasma membrane in ERK7^AID/IAA^ parasites. Instead, micronemes appear haphazardly arranged in the apical parasite cytosol (Figure 5B). Like other reports of conoid defects (Long *et al*., 2017a; Nagayasu *et al*., 2017), we observed functional microneme secretion upon ERK7 depletion and disruption of the apical complex (Figure 3). Our data indicate that micronemes must be secreted through a site distinct from the conoid in ERK7^AID/IAA^ parasites.

**Figure 5:**
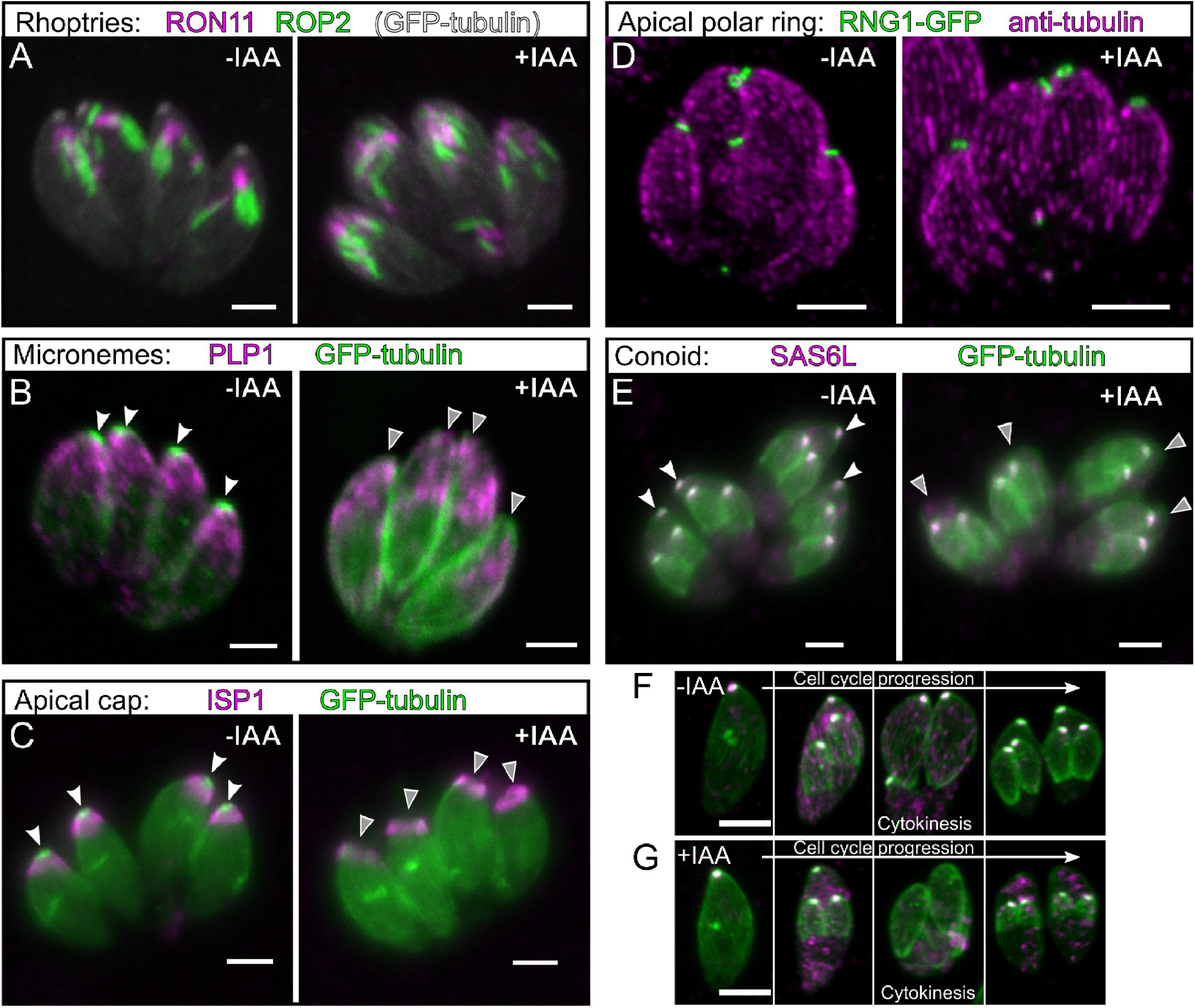
Loss of ERK7 results in disorganized apical structures. (A-E) ERK7^AID^ parasites were grown in +/-IAA and imaged using the indicated markers. All images are maximum intensity projections of confocal stacks. Scale bars: 3 μm. White arrows indicate the apical complex GFP-tubulin foci in mature parasites, which is missing in +IAA parasites (grey arrows). (F-G) ERK7^AID^ parasites were grown for 6h in +/-IAA and imaged (GFP-tubulin – green; SAS6L – magenta). Scale bars: 4 μm.

We also examined markers for distinct substructures in the parasite’s specialized cytoskeleton. The apical cap, as marked by ISP1, appeared to be preserved upon ERK7 degradation (Figure 5C). The apical polar ring marker RNG1-GFP also localized correctly in parasites lacking ERK7 (Figure 5D), consistent with the observed preservation of the cortical microtubules. These data confirm that ERK7 degradation leads to the specific disruption of the parasite conoid.

GFP-tubulin identifies the conoid as bright apical puncta, which was lost upon ERK7 degradation (Figure 5). These data demonstrate that the complete loss of conoid structure observed in the parasite ghosts was not an artifact of sample preparation. We used this phenotype to assess whether ERK7 kinase activity was required for its function in conoid biogenesis. Since we were unable to obtain parasites stably expressing kinase-dead ERK7, we used transient transfection. We transfected ERK7^AID^ parasites stably expressing GFP-tubulin with a plasmid encoding either wild-type or kinase-dead (D136A) HA-tagged ERK7 driven by its native promoter. ERK7^AID^ and ERK7^AID/IAA^ parasites were fixed after 24 h growth, stained with anti-HA, and imaged. As expected, expression of wild-type ERK7 enabled ERK7^AID/IAA^ parasites to form conoids (Supplemental Figure S2D). However, we did not observe rescue of conoid formation in ERK7^AID/IAA^ parasites expressing kinase-dead ERK7 (Supplemental Figure S2D), indicating that kinase activity is required for ERK7 function in conoid biogenesis.

### ERK7 is essential for conoid maturation

To understand the role of ERK7 in apical complex assembly, we tested whether ERK7 degradation altered the localization of other proteins that associate with the conoid. The *Toxoplasma* cartwheel protein ortholog SAS6L is normally associated with the conoidal rings (de Leon *et al*., 2013). SAS6L colocalizes with GFP-tubulin foci in ERK7^AID^ parasites, and, like the conoid-localized tubulin, is lost in ERK7^AID/IAA^ parasites (Figure 5E). Surprisingly, SAS6L and GFP-tubulin foci were found in the growing daughter buds of mature ERK7^AID/IAA^ parasites that had lost their conoids.

Together, our data suggest that in parasites lacking ERK7, apical complex biogenesis initiates correctly but fails catastrophically during its maturation. To identify this point of failure, we infected a host monolayer with ERK7^AID^ parasites expressing GFP-tubulin, and incubated them in IAA for 8 h before fixation. As *Toxoplasma* replicates asynchronously, we could visualize multiple points in the cell cycle in a single experiment. As expected, all ERK7^AID^ parasites had clearly identifiable foci for GFP-tubulin and SAS6L at all points in the cell cycle (Figure 5F). Strikingly, we observed a loss of these foci only in ERK7^AID/IAA^ parasites that had initiated or progressed through cytokinesis (Figure 5G). These conoid-deficient parasites then produced daughters with clear tubulin/SAS6L apical foci. This indicates that without ERK7, the apical complex fails to insert correctly into the mother plasma membrane.

### Conclusions

The cilia-like conoid is a key compartment for secretion and host invasion in *Toxoplasma*, but regulators of its function remain poorly characterized. While ERK7 does not localize to the conoid, its loss-of-function results in the complete and specific disruption of the conoid structure. Our data indicate ERK7 is not a conoid structural protein, yet its loss causes a phenotype much more severe than the disruption of any known conoid-localized protein (Long *et al*., 2017b, 2017a; Nagayasu *et al*., 2017). ERK7 orthologs are early-branching, but understudied, members of the MAPK protein kinase family (Supplemental Figure S1; (Sang *et al*., 2019)), and our data suggest ERK7 is performing a regulatory role in *Toxoplasma* apical complex biogenesis. ERK7 orthologs in model organisms have implicated it in regulating diverse cellular processes, including autophagy (Colecchia *et al*., 2012), protein trafficking and signaling (Zacharogianni *et al*., 2011; Brzostowski *et al*., 2013; Chia *et al*., 2014; Bermingham *et al*., 2017), and ciliogenesis (Miyatake *et al*., 2015; Kazatskaya *et al*., 2017). Remarkably, ERK7 orthologs are found in all Eukaryotic kingdoms, but are expanded in ciliates (Eisen *et al*., 2006) and strikingly missing in organisms that produce no cilia or flagella at any point in development, such as yeast and land plants. We have shown that ERK7 is essential for the biogenesis of a specialized cilium-like structure in *Toxoplasma gondii*, which suggests ERK7 has a conserved role as a critical regulator of ciliogenesis throughout Eukaryota. In metazoan cilia, ERK7 primarily localizes to the basal body (Figure 1A). Apicomplexans lack orthologs for many basal body proteins, and *Toxoplasma* has no identifiable basal body during its asexual stage (Francia *et al*., 2015). We found ERK7 localized to the apical cytoskeletal cap that is delineated apically by the microtubule-organizing apical polar ring and basally by the Centrin-2-containing apical “annuli” (Engelberg *et al*., 2019; Leung *et al*., 2019). We propose that ERK7 functions at the apical cap to regulate the composition as the cytosol transitions to the conoid, akin to ERK7 function at the metazoan cilia transition zone.

## Materials and Methods

### Phylogenetic analysis

Protein sequences for human ERK1, JNK1, p38α, CDK1, ERK5, and ERK7/8, *Toxoplasma* ERK7 (TGME49_233010; ToxoDBv43 http://www.toxodb.org) and the ERK7 kinase domain sequences from the indicated organisms in Figure 1 were retrieved from Uniprot and aligned using Clustal Omega (Sievers *et al*., 2011). The maximum likelihood phylogenetic tree and bootstrap analysis (1000 replicates) were estimated using RAxML v8.1.17 (Stamatakis, 2014).

### PCR and plasmid generation

All PCR was conducted using Phusion polymerase (NEB) using primers listed in Supplemental Table 1. Constructs were assembled using Gibson master mix (NEB). Point mutations were created by the Phusion mutagenesis protocol.

### Parasite culture and transfection

Human foreskin fibroblasts (HFF) were grown in Dulbecco’s modified Eagle’s medium supplemented with 10% fetal bovine serum and 2 mM glutamine. *Toxoplasma* tachyzoites were maintained in confluent monolayers of HFF. ERK7-3xHA and ERK7^AID^ strains were generated by transfecting the RHΔku80Δhxgprt strain (Huynh and Carruthers, 2009) or the same strain expressing OsTIR1 driven by the gra1-promoter with 50 μL of a PCR product using Q5 polymerase (NEB) with 500 bp homology arms flanking a tag (3xHA or AID-3xFLAG) together with 5 μg of a Cas9 plasmid that had been modified to express HXGPRT and also a gRNA targeting the C-terminus of ERK7. The parasites were selected for HXGPRT expression for 2 days and immediately single cell cloned without selection. The ERK7^AID^ strain was complemented by targeting a 3xHA tagged ERK7 driven by its native promoter, together with a bleomycin resistance cassette to the empty Ku80 locus, and selecting with bleomycin. GFP-α-tubulin, actin-chromobody, and SP-mVenus expressing parasites were created by amplifying the FP-marker expression cassette and an adjacent chloramphenicol (or HXGPRT, in the case of SP-mVenus) resistance cassette by PCR and targeting it to a site adjacent Ku80 locus by CRISPR/Cas9-mediated homologous recombination (see Supplemental Table S1) and selecting with chloramphenicol (or MPA/Xan, as appropriate). The original pMIN eGFP-α-tubulin was a kind gift of Ke Hu. The original pMIN actin chromobody-mEmerald was kind gift of Aoife Heaslip. This was vector was modified to replace the mEmerald with TdTomato. RNG1-eGFP, driven by its native promoter, was transiently transfected into parasites which were fixed after 18-24 h growth. Note that ERK7^AID^ parasites developed “escapees” after 1-2 months of passage in which they ceased responding to IAA and ERK7 was no longer degraded. All experiments were therefore conducted with parasites that had been cultured for <1 month after cloning.

### Plaque assays

To measure plaque efficiency, 200 of each parental, ERK7^AID^, and ERK7^AID^ (wt-comp) parasites were allowed to infect confluent HFF in one well of a 6 well plate in the either the presence or absence of IAA. After 7 days, the monolayer was fixed with MeOH, stained with crystal violet, and the resulting plaques counted. All plaque assays were performed in biological triplicate.

### Immunofluorescence

HFF cells were grown on coverslips in 24-well plates until confluent and were infected with parasites. The cells were rinsed twice with phosphate buffered saline (PBS), and were fixed with 4% paraformaldehyde (PFA)/4% sucrose in PBS at room temperature for 15 min. After two washes with PBS, cells were permeabilized with 0.1% Triton-X-100 for 10 min and washed 3x with PBS. After blocking in PBS + 3% BSA for 30 min, cells were incubated in primary antibody in blocking solution overnight at room temperature. Cells were then washed 3x with PBS and incubated with Alexa-fluor conjugated secondary antibodies (Molecular Probes) for 2 h. Cells were then washed 3x with PBS and then mounted with mounting medium containing DAPI (Vector Laboratories). For tyramide-amplification of ERK7-AID-3xFLAG signal, the above protocol was altered as follows. Endogenous peroxidase activity was quenched by incubation of fixed coverslips with 100 mM sodium azide in PBS for 45 min at room temperature. Cells were blocked with 5% horse serum/0.5% Roche Western Blocking Reagent in TBST for 45 min. HRP-conjugated goat anti-mouse secondary antibody (Sigma) was used and tyramide-fluorophore was allowed to react for 30 sec before final washes. Cells were imaged on either a Nikon A1 Laser Scanning Confocal Microscope, a Nikon Ti2E wide-field microscope, or a DeltaVision OMX SIM microscope. SIM stacks were deconvoluted and reconstructed using the manufacturer’s recommended defaults. Primary antibodies used in this study include rat anti-HA (Sigma; 1:500 dilution), mouse m2 anti-FLAG (Sigma; 1:1,000 dilution), rabbit anti-Tg-β-tubulin (1:10,000 dilution), rabbit anti-ROP2 (1:10,000 dilution), mouse-anti PLP1 (1:1,000 dilution; gift of Vern Carruthers), rat anti-RON11 (1:1,000 dilution), mouse anti-ISP1 (1:1000 dilution), mouse anti-SAS6L (1:1000 dilution; final 3 antibodies were gifts of Peter Bradley). Unless otherwise indicated, all micrographs are representative of 20 of 20 images collected. For SIM, images are representative of 5 of 5 stacks per staining condition.

### Western blotting

Proteins were separated by SDS-PAGE and transferred to a PVDF membrane. Membranes were blocked for 1 h in PBS + 5% milk, followed by overnight incubation at 4°C with primary antibody in blocking solution. The next day, membranes were washed 3× with TBST, followed by incubation at room temperature for 1-2 h with HRP-conjugated secondary antibody (Sigma) in blocking buffer. After 3× washes with TBST, western blots were imaged using ECL Plus reagent (Pierce) on a GE ImageQuant LAS4000. Antibodies used in this study include: Rb anti-Tg-β-tubulin (1:1,000 dilution), rat anti-HA (Sigma; 1:500 dilution), mouse m2 anti-FLAG (Sigma; 1:1,000 dilution).

### Invasion and egress assays

For invasion assays, highly infected monolayers were mechanically disrupted by passage through a 27 gauge needle to release them. 3×10^6^ parasites of each strain tested (all expressing GFP-α-tubulin) were added to HFF monolayer grown in a 24 well with coverslip and incubated for 2 h at 37°C. These were then washed 10× with PBS and fixed and prepared for imaging. 3.15 mm^2^ area (~1% of total well) were imaged per experiment and images analyzed in ImageJ. For egress assays, parasites were allowed to grow for 24-30 h in HFF grown in a 24 well with coverslip. Parasites were washed with pre-warmed Hanks Buffered Saline Solution (HBSS) prior to the assay, and then incubated with HBSS containing 1 μM calcium ionophore A23187 (Cayman Chemical) for 10 min at 37°C before fixation and imaging. Live cell egress/PLP1-function assays were conducted as above, using cells plated in thin-bottom 96 wells for fluorescence microscopy (Corning) and imaged in a stage-top environment chamber (Tokai Hit) on a Nikon Ti2E microscope.

### Transmission electron microscopy

Cells were fixed on MatTek dishes with 2.5% (v/v) glutaraldehyde in 0.1M sodium cacodylate buffer. After three rinses in 0.1 M sodium cacodylate buffer, they were post-fixed with 1% osmium tetroxide and 0.8 % K_3_[Fe(CN_6_)] in 0.1 M sodium cacodylate buffer for 1 h at room temperature. Cells were rinsed with water and en bloc stained with 2% aqueous uranyl acetate overnight. After three rinses with water, specimens were dehydrated with increasing concentration of ethanol, infiltrated with Embed-812 resin and polymerized in a 70°C oven overnight. Blocks were sectioned with a diamond knife (Diatome) on a Leica Ultracut UC7 ultramicrotome (Leica Microsystems) and collected onto copper grids, post stained with 2% Uranyl acetate in water and lead citrate. Parasite ghosts were prepared essentially as described in (Nagayasu *et al*., 2017). Briefly, parasites were incubated with 20 μM calcium ionophore in Hanks Buffered Saline Solution at 37°C for 10 min. The parasite suspension was allowed to adhere to a grid, after which membranes were extracted by addition of 0.5%Triton-X-100 for 3-4 min. The samples were then stained with 2% phosphotungstic acid, pH 7.4. All TEM images were acquired on a Tecnai G2 spirit transmission electron microscope (FEI) equipped with a LaB_6_ source at 120 kV. Images in Figure 4A,B of sectioned cells are representative of 14 ERK^AID^ and 22 ERK7^AID/IAA^ vacuoles imaged. Note that 86% of ERK7^AID^ vacuoles imaged had at least one parasite with a visible conoid while 0% of the ERK7^AID/IAA^ vacuoles imaged had any observable conoid structure.

### Figure generation

Data plotting and statistical analyses were conducted in Graphpad Prism v8.3. All error-bars are s.d. and center-values are means. Figure 3E was analyzed by one-way ANOVA with Tukey’s test. All other p-values are derived from two-tailed t-test. All figures were created in Inkscape v0.92.

## Acknowledgements

We thank Josh Beck, Mike Henne, Vasant Muralidharan, and Ben Weaver for helpful comments on the manuscript; the UT Southwestern Electron Microscopy core facility for assistance with sample preparation; and Dorothy Mundy and the UT Southwestern Live Cell Imaging core facility for assistance with SIM image analysis. M.L.R. acknowledges funding from the Welch Foundation (I-1936-20170325) and National Science Foundation (MCB1553334). X.H. was funded, in part, by Cancer Prevention and Research Institute of Texas Training Grant RP160157. M.M. was funded, in part, by a National Science Foundation Graduate Research Fellowship (2019274212).

---

**Supplemental Figure S1:**
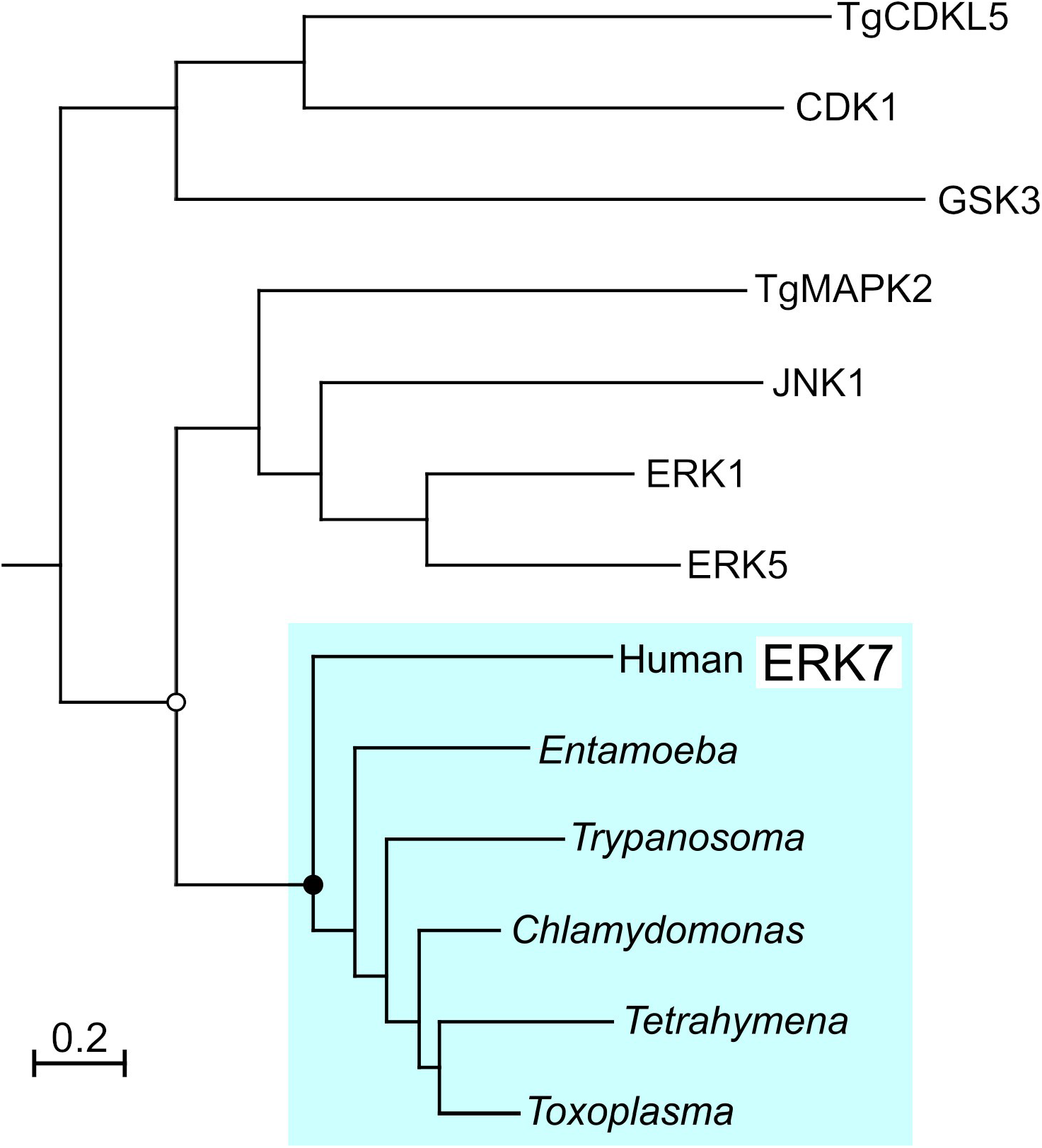
Phylogenetic tree of ERK7 kinase domains from diverse organisms (“ERK8” in human) with *Toxoplasma* CDKL5 and MAPK2 and members of the human CMGC kinase family used as outgroups. Bootstrap values indicated by: black circles: >99%; white circles >90%. Note that ciliates such as *Tetrahymena* have an expanded number of ERK7 paralogs; *Tetrahymena* encodes three distinct ERK7 proteins, though only one is shown here for simplicity. Also note that while *Amoebozoa* such as *Entamoeba* and *Dictyostelium* have apparent ERK7-like kinases, their proteins contain only the kinase domain, and lack the long instrinsically-disordered C-terminus that is present in all other orthologs. Thus, *Amoebozoa* ERK7 likely represent a functionally distinct clade of the ERK7 family.

**Supplemental Figure S2:**
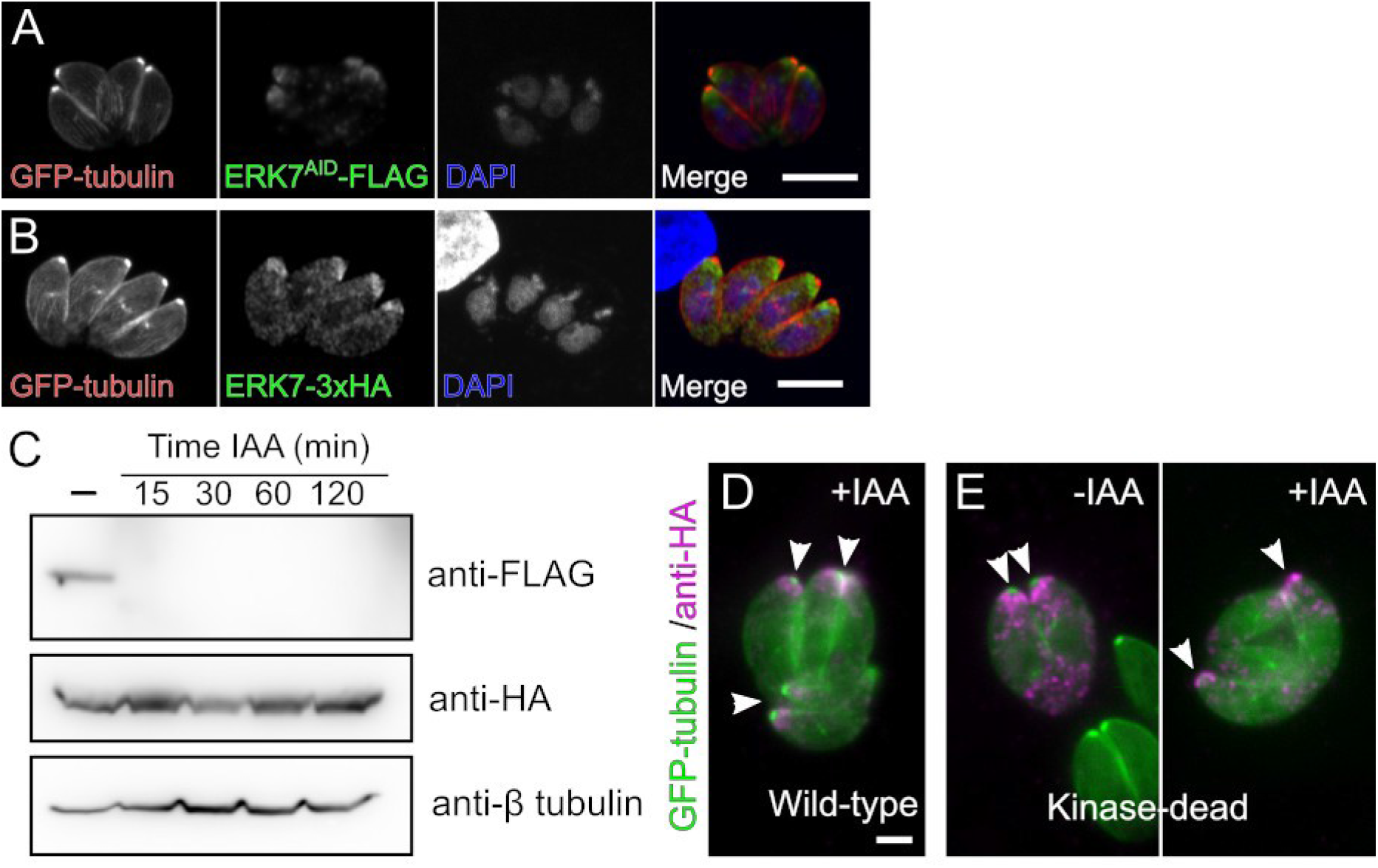
ERK7^AID^ and complementation. (A) ERK7^AID^-3xFLAG (green) was visualized using tyramide enhanced immunofluorescence. GFP-tubulin (red) and DAPI (blue) were used as counterstains. Scale bar 10 μm. (B) An extra, wild-type copy of ERK7-3xHA (green) expressed in the ERK7^AID^ background localizes to the apical cap, just below the GFP-tubulin (red) foci of the conoid. (C) Degradation of ERK7^AID^-3xFLAG does not effect the ERK7-3xHA wild-type copy, as demonstrated by western blot, and the ERK7-3xHA does not effect ERK7^AID^ degradation. ERK7^AID^ parasites were transiently transfected with either (D) wild-type ERK7-3xHA or (E) the kinase-dead mutant D136A and assessed for conoid formation after growth in +IAA media using GFP-tubulin as a marker.

**Supplemental Table 1:**
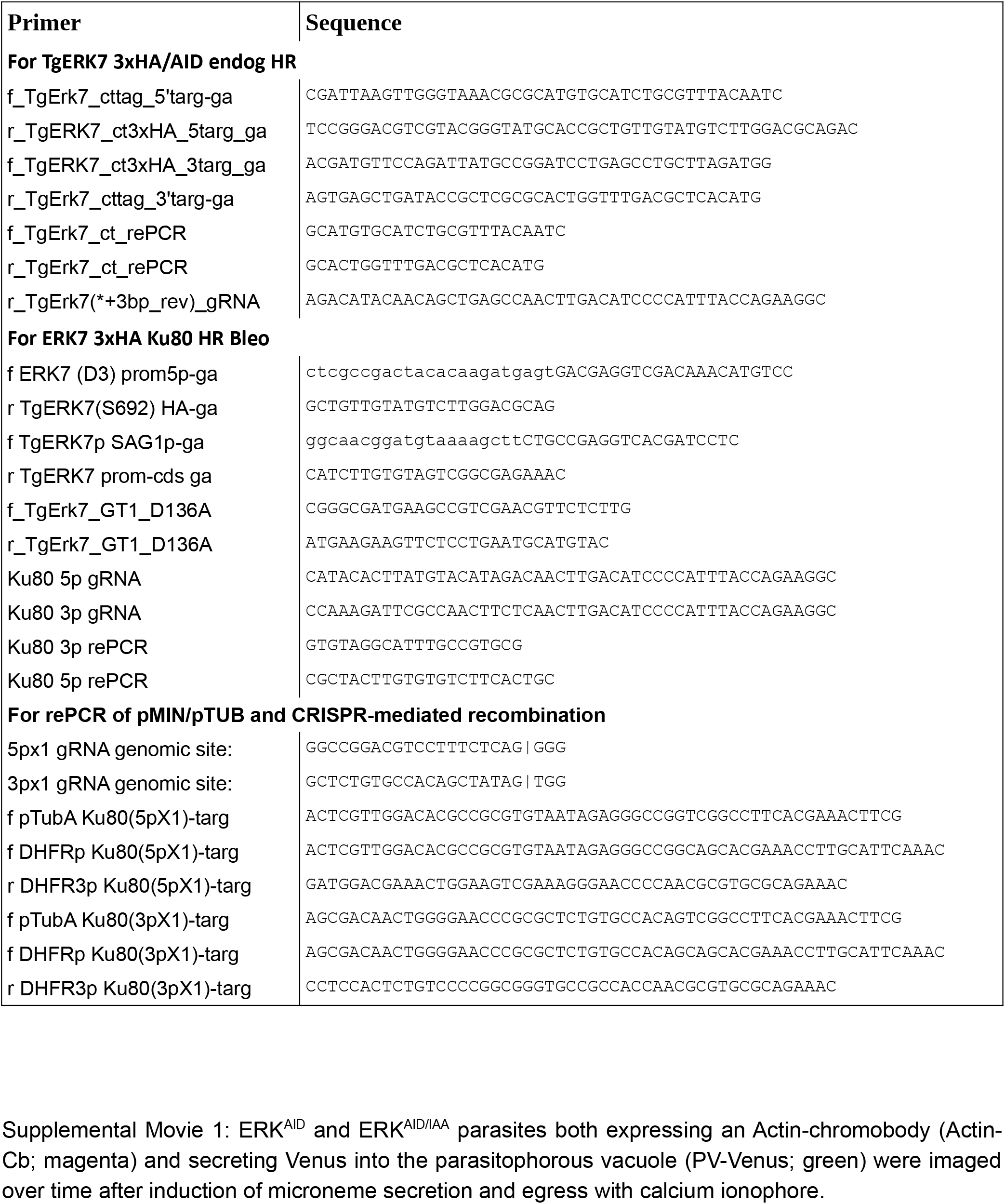
Primers used in this study

Supplemental Movie 1: ERK^AID^ and ERK^AID/IAA^ parasites both expressing an Actin-chromobody (Actin-Cb; magenta) and secreting Venus into the parasitophorous vacuole (PV-Venus; green) were imaged over time after induction of microneme secretion and egress with calcium ionophore.

